# Temperature variability alters the stability and thresholds for collapse of interacting species

**DOI:** 10.1101/2020.05.18.102053

**Authors:** Laura E. Dee, Daniel Okamtoto, Anna Gårdmark, Jose M. Montoya, Steve J. Miller

## Abstract

Temperature variability and extremes can have profound impacts on populations and ecological communities. Predicting impacts of thermal variability poses a challenge because it has both direct physiological effects and indirect effects through species interactions. In addition, differences in thermal performance between predators and prey and non-linear averaging of temperature-dependent performance can result in complex and counterintuitive population dynamics in response to climate change. Yet the combined consequences of these effects remain underexplored. Here, modeling temperature-dependent predator-prey dynamics, we study how changes in temperature variability affect population size, collapse, and stable coexistence of both predator and prey, relative to under constant environments or warming alone. We find that the effects of temperature variation on interacting species can lead to a diversity of outcomes, from predator collapse to stable coexistence, depending on interaction strengths and differences in species’ thermal performance. Temperature variability also alters predictions about population collapse – in some cases allowing predators to persist for longer than predicted when considering warming alone, and in others accelerating collapse. To inform management responses that are robust to future climates with increasing temperature variability and extremes, we need to incorporate the consequences of temperature variation in complex ecosystems.

## Introduction

Climate change is altering climate extremes and variability of environmental conditions [1], yet much of the focus on the impacts of climate change on wild populations remains on how shifts in average conditions will affect dynamics and distributions [2,3]. Yet as average temperatures shift, both temperature variability and the frequency of extreme events, such as marine heat waves, are changing [1,4–6]. Changes in temperature variation and extremes can have profound impacts on individuals and populations, as temperature affects their rates of metabolism, consumption, somatic growth, reproduction, and survival [7,8]. These processes underpin the productivity and resilience of populations and ecosystems [9], on which ecosystem services, such as fisheries yields [10], and population resilience depend. While the potential impacts of temperature variability and extremes have been demonstrated, and in some cases pose a greater risk to species than increases in mean temperature [7,8,11], predicting the importance of temperature variability and extremes for populations, communities, and the ecosystem functions and services they support remains challenging. First, temperature can have both direct and indirect effects that create complex feedbacks and dynamics [12]. Directly, temperature affects species through physiological performance. But temperature can also impact species indirectly, as shown for warming [13,14], by increasing or decreasing important resources or prey species (in turn affecting consumers and predators), changing competitive abilities (altering prey abundance distributions), and altering feeding rates and top-down control (affecting prey) [15–18]. Second, even without shifts in mean temperatures, temperature variability can alter mean vital rates because of non-linear relationships between temperature and processes including growth, reproduction, and mortality (Fig 1; [19]). As a result, the responses of populations to increasing temperature variability and extremes induced by climate change, especially in a community context, remains a key research frontier in marine population dynamics, community ecology, and fisheries science.

**Figure 1.**
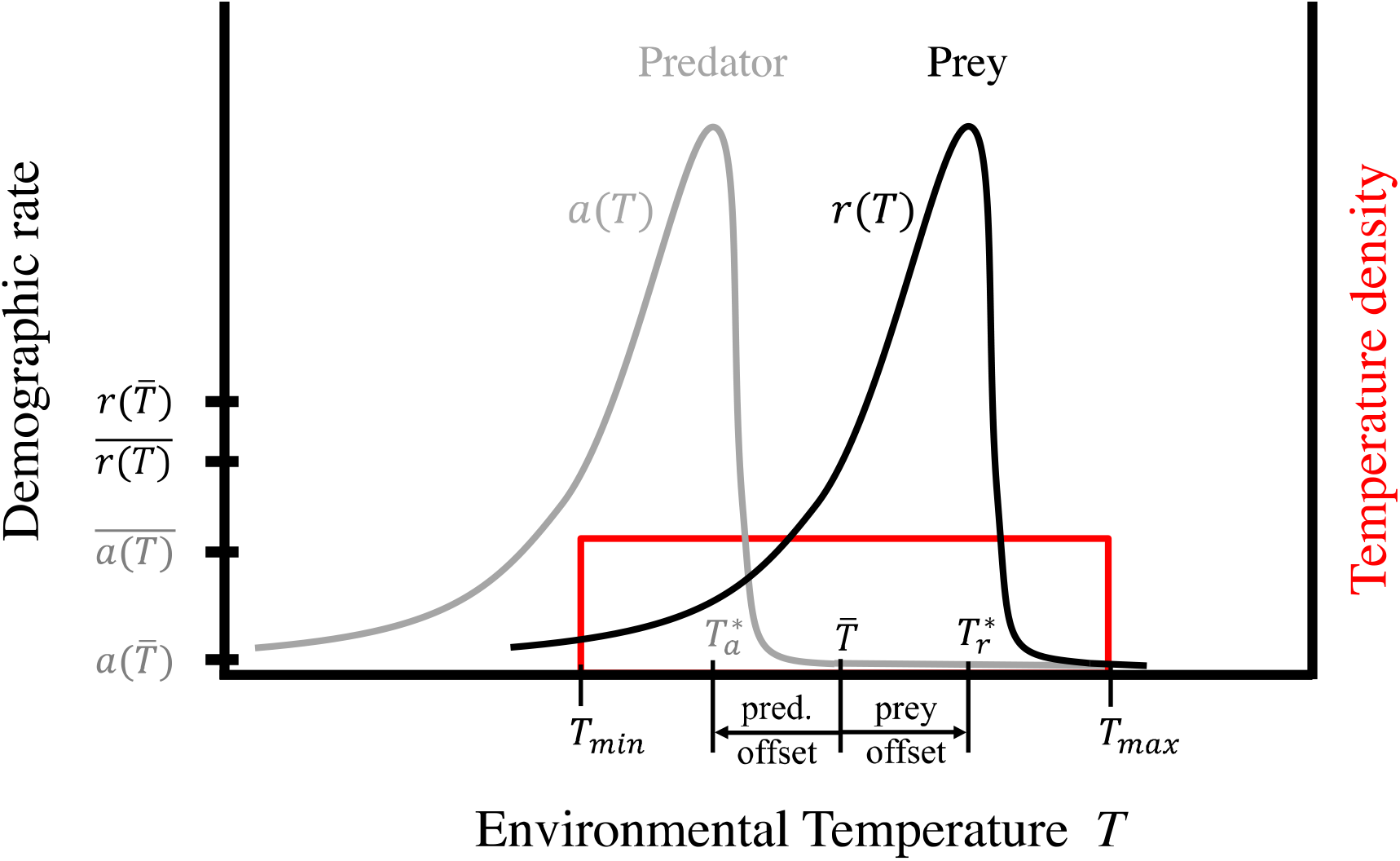
Nonlinear responses to temperature. A conceptual figure of how variable temperatures affect demographic rates, e.g., intrinsic per-capita growth rate *r*(*T*) or attack rate *a*(*T*), following thermal performance curves (TPCs). Each individual of a species has a temperature optimum *T*\* at which its performance is maximized, which may be offset from the mean environmental temperature 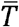. When temperature varies, average demographic rates 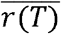 and 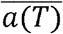 may be higher or lower than demographic rates at the mean temperature 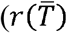 and 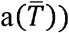 due to Jensen’s inequality. For example, average rates are likely to be smaller for species adapted to their average ecosystem temperature, i.e. 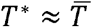. If the range of temperatures encompasses both convex and concave regions of the TPC, the net effect is indeterminate but generally nonzero. In experiments, we vary (1) the amplitude of temperature variability (*T*_max_ − *T*_min_), and (2) how far the TPCs are offset from the environmental mean temperature 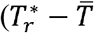 and 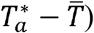. We restrict offsets to be equal in magnitude but have opposite sign, reporting results in terms of the predator’s TPC offset 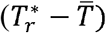.

Directly, temperature variability can impact populations of species in a variety of ways, including by altering the rates of fundamental processes that determine population size, extinction risk, and productivity. Increases in the frequency or duration of exposure to extreme temperatures can induce physiological shock leading to depressed somatic growth or lower survival [20,21]. Temperature variability and extremes above cold or warm events, in contrast to mean temperature conditions, can also alter environmental cues that induce or suppress reproductive cycles leading to skip spawning [22], development [23,24], or hatching [23] that change the number of offspring (i.e. recruitment in fish) or cause reproductive failure. Theory and empirical work shows species’ performance responds non-linearly and asymmetrically to environmental temperature, including intrinsic growth rates and other fitness proxies. This relationship is known as a thermal performance curve (TPC) [8,25,26]. Because of this non-linearity, changes in temperature variance can affect demographic rates differently than changes in mean temperature alone, due to non-linear averaging and Jensen’s inequality (i.e., that the mean of a concave function is smaller than the concave function of a mean, and vice versa for convex functions, see Fig. 1; put in an ecological context *reviewed in* [19]). This also means that if species are well-adapted to the mean conditions, such that their thermal optimum is close to the average environmental temperature (i.e., in the concave part of the thermal performance curve), then increases in temperature variation around the mean are predicted to reduce a species’ performance (e.g., growth rate) [20,27,28]. These impacts scale from individuals to population growth rates [29,30]. Similarly, nonlinear TPCs suggest that higher frequency or duration of exposure to temperature extremes, rather than longer-term temperature averages, shape growth and mortality [31] due to species’ asymmetric responses to higher temperatures and extremes [20]. Collectively, these nonlinear responses and intermittent exposures to thermal extremes suggest that temperature variability can alter population dynamics in complex ways not explained by warming of average temperatures alone [7,30,32].

Simultaneously, temperature can indirectly affect populations through species interactions, thus community and ecosystem responses to temperature variability are complex to predict. Species interactions mediate how temperature impacts a population’s growth, biomass, and dynamics. Because species that interact can respond differently to temperature [33,34], species-specific thermal performances can alter interspecific interactions and resulting population dynamics. For instance, if temperature increases prey growth while predators net growth increase less (e.g., due to increased metabolism for larger bodied-species,[35,36]), prey can outgrow predation windows raising survival [37]. This suggests that if prey thrive directly in response to temperature variation when predators are heat suppressed, prey may flourish in response to heatwaves, and *vice versa.* Understanding the consequences of temperature variability for marine communities therefore requires considering species interactions, because some species benefit from variable temperatures while others lose (Fig. 1).

However, the effects of both temperature variability (or extremes) and species interactions are rarely accounted for in studies of how climate change impacts community dynamics and ecosystem functioning. On one hand, lab studies that quantify the impacts of temperature variability on individuals and populations often focus on the direct physiological effects on performance (e.g., [8,30,32]), with few considering both variation and interactions (*but see* [7,38,39]). On the other hand, most experimental and theoretical studies investigating warming effects on species interactions and their indirect effects do so under different levels of average temperature conditions (e.g.,[15,37]), with less emphasis on variability and extremes (*but see* e.g., [7]). However, when demographic rates depend non-linearly on temperature and interacting species differ in their thermal responses, we anticipate that the net effect of these processes can lead to unexpected outcomes for population dynamics and stability. Indeed, the net effect of temperature variability could be more positive or negative than considering either effect in isolation.

In this paper, we examine how different temperature regimes impact the dynamics of interacting predators and prey, with a focus on multiple types of temperature variability, including increases in temperature variability associated with climate change. We theoretically investigate when considering the combined direct and indirect effects of temperature variability (“net effect”) alters predictions for population productivity, stability, and trajectories through time, relative to considering a constant environment and/or warming average temperatures alone. Specifically, we ask:

1. How does increasing temperature variability affect population size through time, occurrence of collapse, and stable coexistence of both predator and prey, relative to a constant environment?
2. How do these effects depend on the predator’s and prey’s TPCs relative to the temperature variability regime, and relative to one another?
3. What are the net *versus* direct effects of temperature variability on these properties when species interact, and when are direct versus net effects acting in opposite directions?
4. How do effects of temperature variability compare to and/or modify the effects of increasing mean temperatures?

We hypothesize that the extent to which the effects of temperature variability shift predictions about population growth, size, and stable coexistence beyond warming average conditions will depend on the 1) thermal variability regime, 2) strength in species interactions, and 3) overlap in TPCs of interacting species (Fig. 1). We motivate our theoretical investigation with a marine predator-prey system, though the model applies more generally to predator-prey systems experiencing both temperature variability and rising temperatures.

## Methods

### Model

We model a predator-prey system in which a subset of key parameters depend upon temperature. In line with prior investigations of how temperature affects community dynamics [40–43], we use a Rosenzweig-MacArthur model with a prey population *x*(*t*) and predator population *y*(*t*) changing as a function of time *t*:

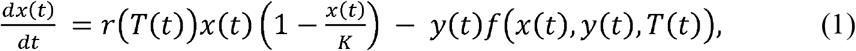

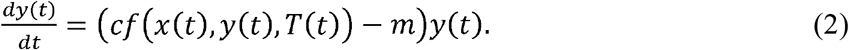

Here, *r* is the temperature-dependent intrinsic growth rate of the prey population, *K* their carrying capacity, *T*(*t*) is temperature at time *t*, *m* the density-independent mortality in the predator, and *f* denotes the Holling type II functional response of the predator:

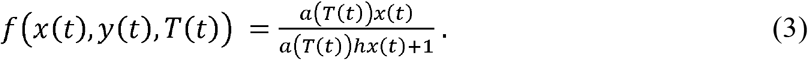

Here *a* denotes attack rate, which depends on temperature; *h* is handling time; and *c* is conversion efficiency.

In the absence of temperature effects, this Rosenzweig-MacArthur model has three equilibria: joint extinction, extinction of the predator with the prey at carrying capacity, and a coexistence equilibrium. The coexistence equilibrium is defined by:

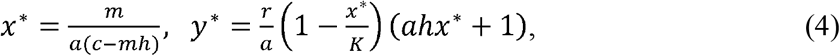

which requires that *aK*(*c* − *mh*) > *m*. We use this equilibrium to set the starting conditions, in terms of the populations levels (abundances), to provide consistent initial conditions under which there are no transitory dynamics in the absence of temperature-dependence.

We focus on understanding how temperature variability influences population dynamics via impacts to the predator and the prey species. Particularly, we model how both the attack rate (*a*) of the predator [44] and the intrinsic growth (*r*) of the prey [8] depend on temperature. Following current theory and empirical studies, we assume *r*(*T*) and *a*(*T*) follow unimodal non-linear relationships, with maxima at 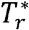 and 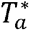, respectively (Fig. 1). Although the shape of thermal performance curves (TPCs) will vary among species, populations, and life stages, TPCs are generally considered to have a steep, negative drop-off in performance above optimal temperatures, but rapid gains in performance at lower temperatures (Fig. 1) [33,45,46]. In our model, we use species-specific asymmetric hump-shaped TPC following Gårdmark et al. (in prep) and [47,48]; see SI for details.

We focus on the temperature-dependence of these two parameters, attack rate *a* and prey growth rate *r*, but recognize that temperature also could influence other parameters in our model (e.g., carrying capacity *K*, handling time *h*, and mortality rate *m*) [44,49]. We choose to examine the temperature dependence of *a* and *r* because these are the parameters for which we have the most unequivocal information (*see* [50]). For example, carrying capacity may also vary with temperature [49], but that parameter is not mechanistic in the Rosenzweig-MacArthur model. That is, *K* may be determined by many factors including primary productivity, nutrient supply, and habitat availability, and temperature may affect each of these in different and complex ways that are more likely linked to exogenous factors that this model does not incorporate.

### Simulations

The way in which temperature variability affects growth and attack rates depends on the distribution of temperatures to which species are exposed and the species’ TPCs. Because TPCs for a prey and its predator are likely to differ (Fig. 1), we explore how overlap of TPCs and their relation to temperature distributions affect population dynamics for both species. Thus, we conduct several simulations that combine and vary both: the temperature regime and how species’ TPCs relate to one another. For each simulation, we examine how the temperature regime influences average temperature-dependent demographic rates (*r* and *a*), long-run population means for both predator and prey, and the equilibrium type (either extinction or coexistence with either fixed densities or cycling dynamics). Here, we define ‘*long-run’* predator and prey populations based on the last 10% of the simulated time steps (final 200 out of the 2000 time steps), which provides time required to reach a stationary distribution (Fig. S1), if one exists.

## Temperature regimes

### Scenarios of temperature regimes for means and variability

The temperature regimes we consider are (1) temperature variability of different amplitudes, but constant mean temperature (‘variability-only’), (2) increasing mean temperature, but no variability (‘warming-only’), (3) increasing mean temperatures and constant temperature variability of different amplitudes (‘warming-and-constant variability’) and (4) both increasing mean temperatures and increasing temperature variability (‘warming-and-increasing variability’).

We generate the temperature variability and warming regimes as follows. In the variability-only regime, temperatures oscillate linearly between a maximum *T*_max_ and minimum *T*_min_, completing one cycle per time period (see SI) to reflect seasonality. In this scenario, increases in temperature variability do not affect the mean temperature 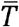. In the warming-only regime, temperatures increase linearly from *T*_min_ to *T*_max_ over the course of the simulation. To facilitate comparison, all warming-only scenarios share the same 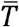, such that the variability-only and warming-only regimes have the same overall distribution of temperatures for a given *T*_min_ and *T*_max_. We generate these regimes for different sets of *T*_min_ and *T*_max_, which we refer to as “amplitudes.” Finally, the warming-and-variability scenarios combine the linear warming and linear oscillation effects with different amplitudes, where both warming and variability share a specified amplitude (see SI).

### Scenarios with stochastic events driving temperature variability

In addition to the deterministic temperature changes in the form of seasonal variation, warming, or both, <u>we also simulate temperature variability that is stochastic, to test the sensitivity of our results.</u> We examine two types of stochastic temperature changes (see SI section 1.2). First, we examine major “events” modeled after El Niño; when such an event occurs, the temperature shifts up during a given year. Second, we examine smoother stochastic variability in the form of a Gaussian process with squared exponential covariance function, allowing for both variance and autocorrelation to increase through time. In both cases, we add these stochastic temperature deviations on top of our baseline scenario of oscillating (seasonal) temperatures - enabling us to focus solely on the two types of variability rather than the combination of variability and warming.

### Examining how predator-prey TPC overlap and trophic structure modulates effects of temperature

To investigate the extent that TPC overlap between predator and prey species determines the impact of temperature on each population and their coexistence, we systematically vary the extent of TPC overlap between predator and prey species for each temperature scenario. For each temperature regime, we consider a range of configurations of species’ TPCs, defined by the offset 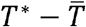 (in degrees Celsius) between each species’ optimal temperature and the mean environmental temperature (see Fig. 1). Specifically, we consider cases where the TPCs for the predator and prey have offsets of the same magnitude but opposite sign, so that the location of both TPCs is defined in terms of the prey offset (as in Fig 1). When that offset is zero, the predator and prey TPCs overlap exactly. Table S1 defines parameters for these simulations.

Finally, we conduct an additional set of analyses in which we introduce a second prey species with different thermal affinities than the first prey. To do so, we use a multispecies Holling type II functional response [51], to maintain the same functional form assumptions as above, in order to isolate the effect of adding an additional prey on stable coexistence. To test the influence of multiple prey species with different TPCs on co-existence of predator and prey species, we shifted the first prey TPC and the predator TPC in opposite directions as before, but fixed the second prey TPC at the environmental mean, to facilitate comparison with the single prey case.

### Calculating direct effects versus net effects of temperature via species interactions

To quantify the importance of accounting for species interactions when studying the effects of temperature variability, we contrast the direct and net effects (direct plus indirect) of temperature variability on the predator population. For each combination of temperature regime and species’ TPCs, we track the predator population for three simulations, with growth and attack rates set to (*i*) 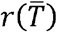 and 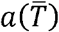; (<u>*ii*</u>) 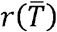 and *a*(*T*); and (*iii*) *r*(*T*) and *a*(*T*); rates defined either at the actual temperature *T* or at the mean temperature 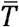 ignoring variability. Thus, simulation (1) ignores all effects of temperature variability, simulation (2) ignores the effects of variability on prey growth, and simulation (3) examines the effects of temperature variability on both species. We define the net effect of temperature variability as the difference in long-run predator population between simulations (3) and (1), while the direct effect is defined by the difference between simulations (2) and (1).

## Results

### Effects of temperature variability on long-run populations

Increases in temperature variability produce either stable coexistence, predator-prey cycles, or extinctions (Fig. 2), depending on the difference in predator and prey TPCs and parameters relating to species interactions (predator conversion efficiency and attack rate) (Figure 1; Figure S1). When TPCs for predator and prey coincide (i.e., are identical), variable temperatures can drive the predator extinct from an equilibrium that is stable under conditions without temperature variability (Figure 2b; Fig S1). For higher parameter values of predator conversion efficiency (*c* = 0.3 in Fig. 2a versus *c* = 0.1 in Fig. 2b), however, increasing temperature variability can stabilize the system, causing a shift from cyclic dynamics (limit cycle) to a stable equilibrium point (fixed point) (Fig 2a). The opposite effects can arise when the TPCs for the two species are offset so that the prey’s temperature optimum is higher. Specifically, when peak attack rates are high, variability can save the predator from extinction, if the prey has a much higher optimum temperature than the predator, or destabilize an otherwise stable equilibrium by inducing cyclic dynamics when the difference in TPCs between predator and prey is less (Fig. 2c). Even when variability does alter the type of equilibrium that arises, equilibrium population levels can be driven up or down.

**Figure 2:**
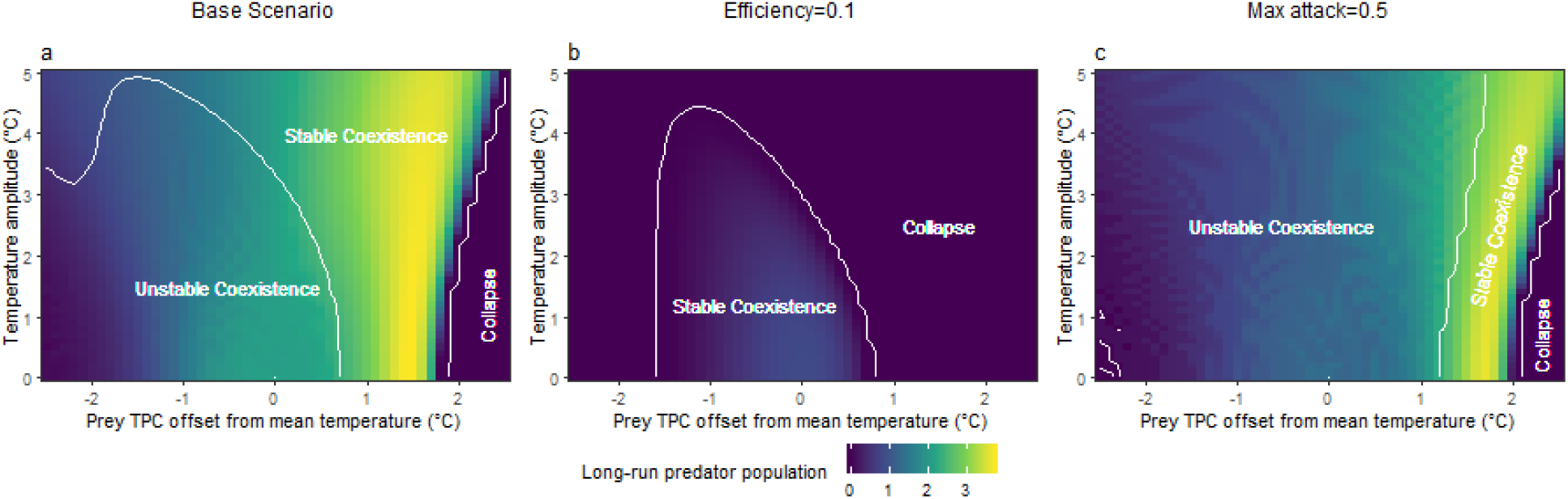
Stable predator collapse or predator-prey coexistence depends on both the offset in the predator and prey thermal performance curves (TPCs) and amplitude of temperature variability. Effects of offset in the predator and prey TPCs (x-axis) and amplitude of temperature variability (y-axis) on predator population abundance (colors) and the type of equilibrium that arises: predator collapse or predator-prey coexistence, and whether the latter is stable or unstable (e.g., cycles or oscillatory behavior). Base scenario parameters (a) are mortality *m* = 0.2, carrying capacity *K* = 20, conversion efficiency *c* = 0.3, maximum attack rate *a* = 0.3, and handling time *h* = 0.3, whereas species interactions are modified in (b-c) by lowering conversion efficiency *c* = 0.1 (B) or increasing attack rate *a* = 0.5 (C). TPC parameters are in Table S1.

The presence of a second prey species with different thermal affinities from the first prey increases the parameter space with stable coexistence (Fig. S5). Predator feeding on an alternative prey with TPC at mean temperature stabilizes dynamics and prevents extinction, especially when the original prey has a much lower optimal temperature than its predator (negative prey TPC offsets, cf. Fig. 2 and Fig. S5) and for predators with low conversion efficiency (Fig. S5).

### Direct versus net effects of temperature variation

Including indirect effects can qualitatively change the impact of temperature variability on populations (Fig. 3; Fig. S2). The true, ‘net’, effect on long-run population levels can differ from that suggested by studying the direct effect alone, both in terms of direction of the response (Fig. 3) and effects on the stability of predator-prey dynamics (Fig. S2). In other cases, the direct effect of temperature variability may imply an unstable equilibrium, but after accounting for indirect effects, the equilibrium remains stable (Fig. S2).

**Figure 3:**
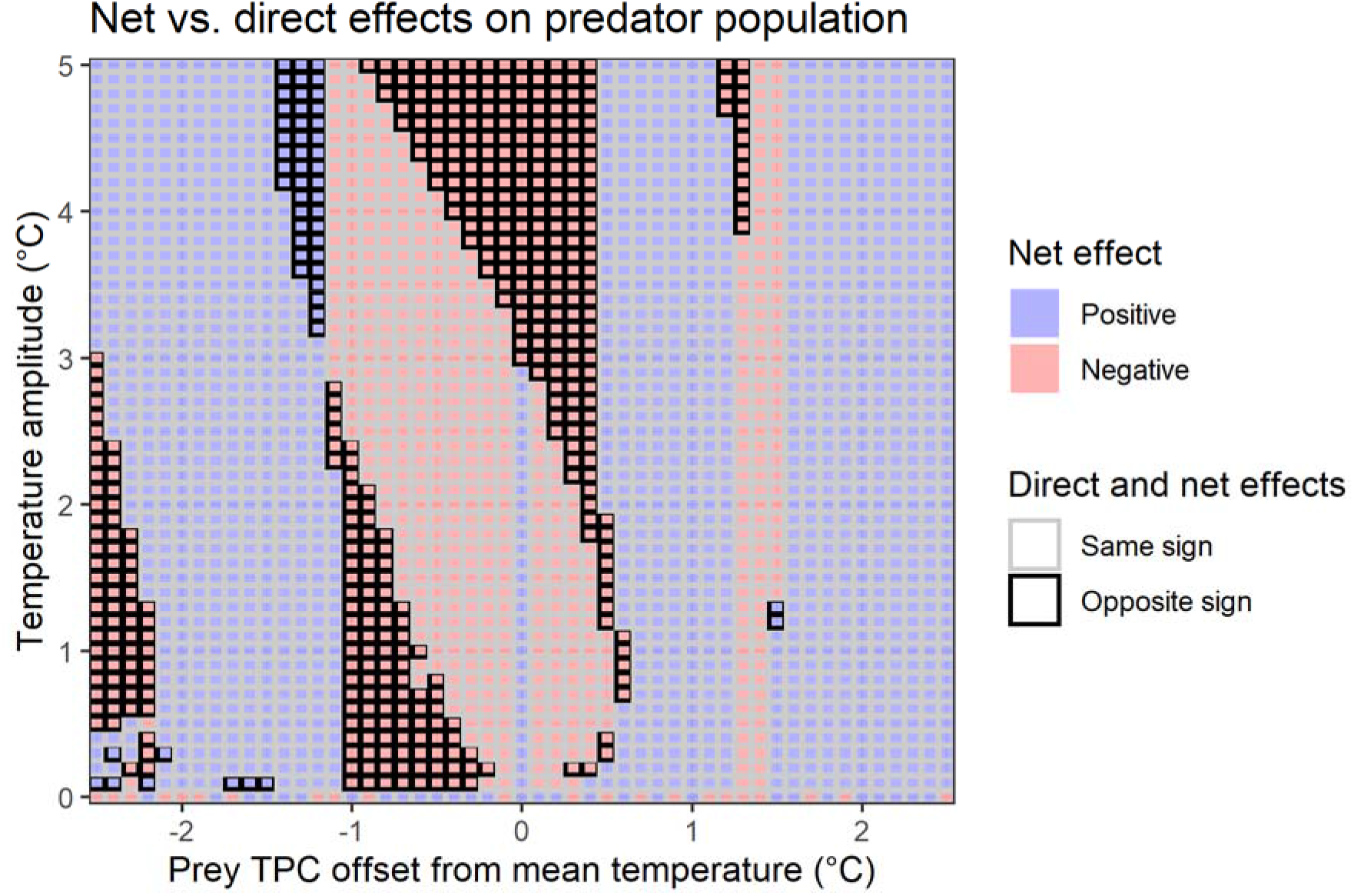
The net and direct effects of temperature variability on predator population. Net effects (blue: positive; red: negative) of temperature variability on long-run population levels for the predator, as a function of how far the prey TPC is offset relative to the mean temperature (x-axis) and the amplitude of temperature variability (y-axis). Outlines indicate whether net effects have the same sign (black outline) or not (light gray outline) as when only considering the effects of temperature variability on the predator, ignoring the temperature-dependence of its prey (‘direct effects’). For example, red-filled and black-outlined regions indicate parameterizations where considering only temperature effects on the predator would suggest a positive effect of temperature variability for the predator population when the true net effect is negative, due to species interactions. Parameters as in Fig. 2a with TPC parameters in Table S1; see Fig S4 for results from additional parameterizations.

### Effects of increased temperature variability versus increased temperature mean

The effects of increasing mean temperature on long-run population sizes without shorter-term variability are qualitatively different than those of variability without warming, even for an identical distribution of temperatures experienced during the simulation (Fig. 4). For example, increasing mean temperature on predator and prey with completely overlapping TPCs can drive the predator extinct (Fig. 4, *red lines*) as higher temperatures eventually lead to extended periods of low attack rates (which asymptote towards 0). In contrast, variability with the same distribution of temperatures can lead to a stable coexistence (Fig. 4, *blue lines*). When mean temperatures increase and variability is present — whether constant (Fig 4, *purple lines*) or increasing (Fig 4, *orange lines*) — the predator population survives for longer than under increasing mean temperature alone.

**Figure 4.**
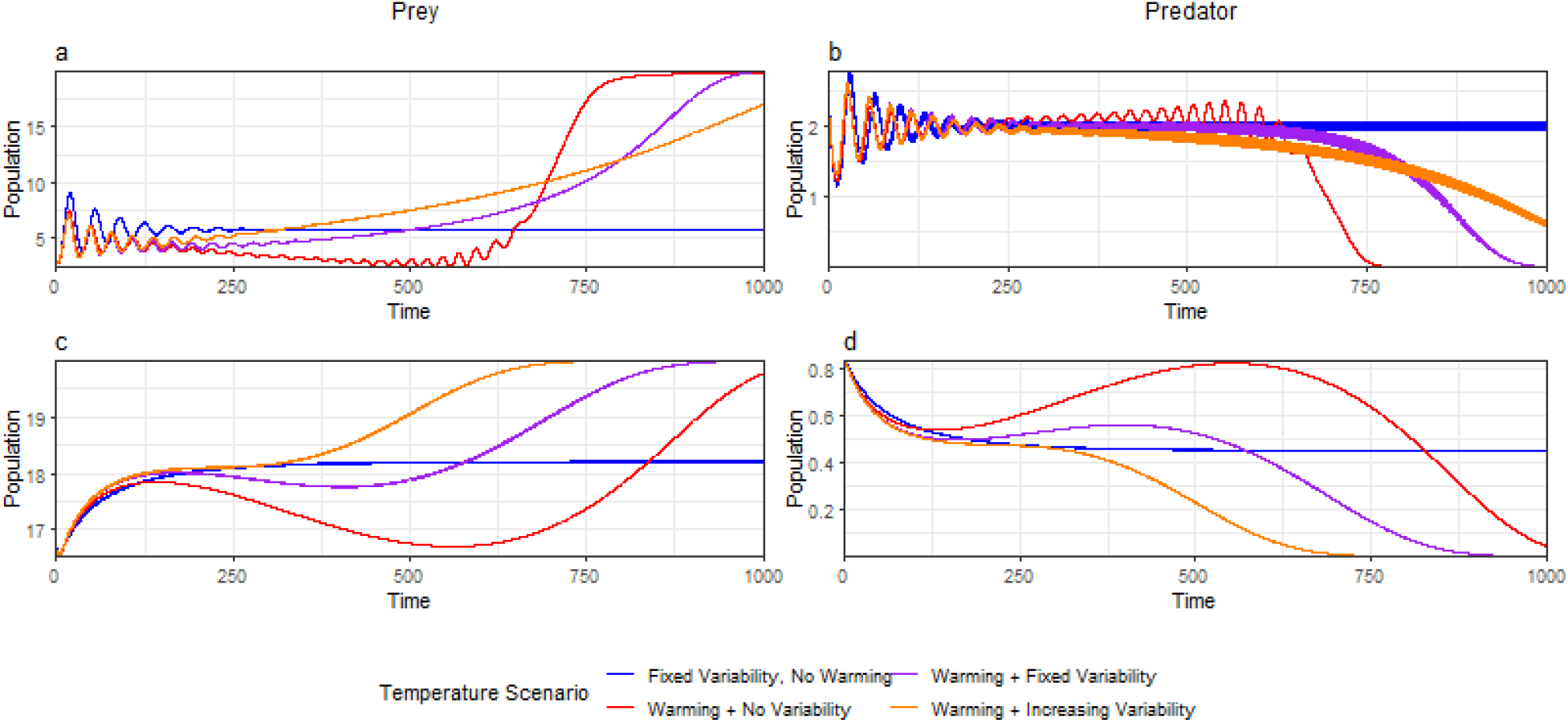
Interacting effects of warming and temperature variability. Population trajectories for the prey (a,c) and predator (b,d) under scenarios of warming (increasing mean temperatures) with no variability (red), constant temperature variability with constant mean temperatures (blue), increases in mean temperatures with constant variability (purple), and increases in both mean temperatures and temperatures variability (orange). Variability on top of warming can either delay (panel b) or accelerate (panel d) predator collapse. Panels a and b reflect model parameters as in Fig 2a with *T*_max_ − *T*_min_ 8°*C*; panels c and d reflect model parameters as in Fig 2b with *T*_max_ − *T*_min_ 2°*C* TPC parameters are in Table S1 with zero offset.

Accounting for stochastic variation in temperature does not change our qualitative conclusions (Figure S6). While moderate stochastic temperature changes knock population trajectories out of asymptotic convergence to equilibrium levels, they do not qualitatively change the means of the long-run population trajectories in response to temperature (Fig. S6)

## Discussion

Increases in temperature variability can influence populations of interacting species in ways not predicted by considering increases in mean temperatures or the direct physiological effect of temperature variation alone. First, a focus on mean temperature alone misses the highly non-linear responses of species’ demographic rates to changes in temperature [52]. Temperature variability, by directly creating complex nonlinear demographic responses, can alter both the viability and stability of populations in either direction (Fig. 2). Second, the net effect of temperature variability when species interact can 4differ from that predicted by the direct effect of temperature on a single species, and even be of opposite sign (Fig. 3). Thus, ignoring indirect effects and making predictions about population dynamics solely from individual demographic rates (e.g. growth) may create erroneous expectations, somethings in the opposite direction (Fig. 3; Fig. S2). This highlights the importance of considering temperature effects in contexts with both intra- and inter-specific interactions. Third, results depend on the differences in predator and prey TPCs (i.e., if the predator TPC is optimized at a higher, identical, or lower temperature than its prey). Finally, results from our model show ignoring temperature variability could over-predict negative impacts of warming on population and community trajectories. Even under warming temperatures, temperature variability results in periods in which temperatures return to a range under which species can grow (Fig. 4), though our results show that the predator populations may still eventually collapse as warming intensifies (Fig. 4d). These findings have important implications for natural communities as temperature variability is predicted to increase further due to global warming.

That temperature variability increases species persistence can also be seen in single species models, which provide insights into when this ‘rescue effect’ occurs. This ‘rescue effect’ depends on how temperature variability influences temperature-dependent demographic rates for hump-shaped TPCs. In a single-species logistic model with temperature-dependent growth, the prey population at time *t* depends only on the average growth rate 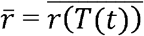 of that time period [53]. Depending on the curvature of the TPC over the range of temperatures that a species experiences, temperature variability can either raise (in convex parts of the TPC) or lower average growth (in concave parts of the TPC), with concomitant change in population levels (Fig. 1; *see also* [54, 55]). If warming drives the mean temperature above the prey’s optimal temperature, then temperature variation will span a convex portion of the species TPC, thereby raising average growth and prolonging population persistence (as seen in Fig. 4).

Similar insights can be gleaned about the effects of variability on average rates for interacting predator and prey species. Temperature variability is likely to lower average attack rates when the temperature distribution is centered near the peak of the predator’s TPC (Fig. 2 when TPC offset is around 0), whereas it results in higher average attack rates at lower temperatures (Fig. 2 when TPC offset is large and positive), such that temperature variation occurs in the convex part of the predator’s TPC. Temperature variability in these ranges of the TPC thus promotes predator persistence by resulting in sufficiently high attack rates on average. In fact, simulations using the average predator attack and prey growth rates that arise under variable temperatures yield similar overall patterns of stability and coexistence (see Fig. S3 and SI section S1.3 for details).

Importantly, our results demonstrate that understanding how a particular temperature regime affects predator populations requires knowledge of temperature dependencies of *both* species and how they interact. The true ‘net’ effect of temperature variability on predator populations may be positive or negative; moreover, because of indirect effects, such responses may differ in sign from predictions made based on changes in attack rate (or other properties in a single species) in isolation (Fig 3). The potential for indirect effects on equilibrium populations is evident from equation (4): prey levels are inversely related to attack rates, while predator levels are proportional to prey growth rates.

The difference in TPCs of interacting species dramatically alters the response of ecological communities to variation in temperature (Fig. 2 & 3). Large differences in temperature optima from 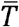 lead to predator extinction, regardless of which species has the higher optimum temperature. If prey have lower temperature optima, predator extinction occurs at even lower levels of temperature variability for the same magnitude of TPC offset, compared to if the predator has the lower temperature optimum. The rapid decline of the TPCs above a species’ optimum explains this result. Average growth for the prey (or attack rate for predator) is substantially lower when temperature vary above that species’ optimum temperature compared to below (i.e. if the temperature optima is shifted downward rather than upward by the same amount). If the prey has a lower temperature optimum than the predator, high temperature variability results in prey population depletion by increasing predator attack rates much more than prey growth rates, eventually leading to predator extinction. These results are consistent with findings from other contexts showing that imbalances in temperature sensitivities of species metabolism and ingestion have important consequences for community dynamics. For example, for invertebrates, metabolism increases faster than ingestion [56,57], which can lead to starvation, and theoretical analyses indicate that this increases population stability but also the risk of starvation and extinction of predators [13,40].

Comparing the effects of temperature variability with those of warming mean temperatures reveals their different impacts on species coexistence and stability, even when the distribution of temperatures under the two regimes is identical. When temperatures warm, long-run predator populations suffer much more than if the same magnitude of temperature change is experienced in a cyclical fashion (Fig. 4). Warming eventually drives predator attack rates to low levels, leading to population declines from natural mortality and a predator population that is not viable. Prey growth also slows, but with reduced predation, the prey population persists. In contrast, under a variability-only scenario, periods of low attack rates are intermixed with stretches of higher attack rates, such that the predator population can survive. Consistent with that intuition, adding variability to a temperature regime with warming can prolong the existence of the predator population, though in other parameterizations variability can accelerate collapse.

We designed our models to quantify the effects that temperature variability can impose on interacting species. However, these models include several simplifying assumptions. Our models assume only the net population growth rate of prey and the attack rate for predators are affected by temperature, but carrying capacities [58], conversion efficiencies [59], mortalities [60,61], and handling times [44,60,62] may also depend on temperature. We also include thermal sensitivity in phenomenological rather than explicitly mechanistic forms. More realistic models (e.g., size-structured, age-structured, or individual-based models) could parse out how maturation, fecundity, mortality, individual growth, and consumption simultaneously respond to thermal regimes in different individuals or size-classes when species interact. For example, accounting for both the temperature-dependent processes underlying body growth and for within-population size structure have proven important for understanding how both consumer populations [35] and food chains [63] vary with temperature. However, while this has been addressed in studies of warming of mean temperatures [35,63,64], it remains to be done for increasing temperature variability. Both the complex responses to increasing temperature variability and the importance of accounting also for indirect temperature effects via interacting species that we demonstrate suggest that predicting how temperature variability would impact size-structured food webs cannot be done *a priori*. Our study thus calls for addressing temperature variability effects in food webs with approaches accounting for bioenergetics processes and the size-dependence of species interactions.

We demonstrate that the presence as well as the form of temperature variability influence species persistence and coexistence and interact with the effects of warming mean temperatures. Future work could consider the following additional complexities in predicting the effects of temperature dynamics on interacting species in particular systems. First, thermal variability with autocorrelation (e.g., persistent heatwaves) can induce not only prolonged shifts in growth but also mortality, even leading to mass mortality, for instance from oxidative stress shaped by temperature maxima rather than mean temperature [21]. Mass mortality events may be induced by short-term exposure to high temperature, and the threshold temperature for mortality can decrease with greater exposure duration [65], whereas high-frequency variability can also reduce negative effects, such as coral bleaching [66]. Second, the impacts of these short-term events are likely to depend on the species’ generation time, relative to time scale of the perturbation. For instance, short-lived species may suffer high mortalities if exposed to unfavorable climate conditions occurring during its short lifespan, compared to longer-lived organism that may better buffer against short-term events. Third, acclimatization [67,68], or short-term evolutionary responses [48,69] to temperature changes could result in inaccurate predictions from models based on historical observations or experiments that are conditional on the environmental history or genotypes. Nevertheless, our results show temperature variability can alter predictions compared to accounting for increases in temperature means alone, indicating the need for considering temperature variability shapes population stability, collapse, and coexistence when species interact (Fig. 2; Fig. 4).

Finally, climate change in the sea is more than warming, variability, and frequency of extreme temperatures. It also encapsulates changes in dissolved oxygen, pH, surface irradiance, salinity, and circulation dynamics -- biophysical changes that are often correlated with temperature fluctuations (IPCC 2019). We chose to focus on temperature dynamics for several reasons. First, there has been a long history of documenting short and long-term variation in ocean temperature either *in situ* or using satellite reconstruction. In contrast, for other biophysical changes, such widespread and high-resolution data collection and reconstructions are more isolated, sporadic or only recently developed. Second, temperature impacts on physiology and population dynamics have long been a focus in fisheries ecology. Thus, we focus on variation in temperature to quantify how variability in abiotic stressors can alter dynamics with the acknowledgment that other stressors and variability therein are also central in shaping populations and communities and may exhibit independent and multiplicative stresses on communities in unpredictable ways. Here we show that the inherent complex direct and indirect responses of populations and communities to gradual linear temperature changes that characterizes climate warming versus the non-linear and extreme changes that characterize climate variability is challenging. This challenge requires that, before including additional and important complexities, we deeply understand the interplay of climate warming and variability.

## Conclusions

Climate change is increasing not only mean temperatures but also temperature variability (IPCC 2019). Understanding the consequences of temperature variability for population trajectories and dynamics is critical for anticipating how climate change will affect the productivity and stability of animal communities that support important functions and services. Here we find that shifts in temperature variability can destabilize, stabilize or lead to predator collapses, depending on interaction strengths and differences in thermal performance between predators and prey. Our results also show that impacts on species’ growth from concurrent changes in variability with warming can change predictions from considering warming alone. Counterintuitively, temperature variability can help a population that would otherwise go extinct due to warming, when warming negatively impacts population growth, and some forms of variability can offset these effects. In other cases, however, ignoring increases in temperature variability associated with climate change leads to underestimation of predator extinction risks.

Our results contribute to a growing understanding of how temperature variation will alter life in the oceans, though the theoretical results extend more generally to other predator-prey systems facing variability. Our findings call for future studies advancing the theory on increasing temperature variation in foodwebs. In particular, we encourage accounting for within-species structure and variation in TPCs, as well as testing this in experimental studies in interacting species (over temperature ranges large enough for the non-linear responses to matter). Moving beyond a focus on mean temperatures alone, to advance our understanding of the consequences of temperature variation in complex ecosystems, can improve our ability to inform management responses that are robust to future climates with increasing variability and extremes.

## Acknowledgements

We thank B. Kendall, D. Bradley, R. Gentry, and C. Costello for discussions that benefited earlier iterations of this research, and the Dee lab and J. Ashander for feedback on this draft. This work was partly supported by the Swedish Research Council (Vetenskapsrådet, grant 201503752 to AG), and by the FRAGCLIM Consolidator Grant, funded by the European Research Council under the EU Horizon 2020 research and innovation programme (Grant 726176 to JMM).

