## Supplemental Materials for "Temperature variability alters the stability and thresholds for collapse of interacting species"

**S1. Additional detail on methods**

***S1.1 Thermal Performance Curves***

We used an asymmetric hump-shaped temperature dependence curve that scales the rates in relation to their value at a reference temperature $T_{ref}$. That curve is based on the Sharpe-Schoolfield relationship [1,2], reformulated in relation to a known rate ($p\left( T_{ref} \right)$) at a reference temperature$T_{ref}$ (Gårdmark et al., in prep.). In this formulation, the rate $p\in\{r,a\}$ at a given temperature $T$ is obtained from:

$$p\left( T \right)=p\left( T_{ref} \right)e^{-\frac{E_{A}}{k}\left( \frac{1}{T}-\frac{1}{T_{ref}} \right)}\cdot\left( {1+e}^{-\frac{E_{D}}{k}\left( \frac{1}{T}-\frac{1}{T_{D}} \right)} \right)^{-1}\cdot\left( {1+e}^{-\frac{E_{D}}{k}\left( \frac{1}{T_{ref}}-\frac{1}{T_{D}} \right)} \right).$$

Here, $E_{A}$ is the activation energy of the rate (governing the increase in the rate of temperature), $E_{D}$ the deactivation energy of the rate, $T_{D}$ is the temperature at which the rate has declined to half of its maximum value, and, and $k$ is the Boltzmann constant. The parameters $E_{A}$, $E_{D}$, $T_{D}$, $T_{ref}$, and $p(T_{ref})$ are specific to the parameter *p*; we omit further subscripts for brevity. The optimum temperature $T^{*}$ for this TPC is at

$$T^{*}=\frac{E_{D}T_{D}}{E_{D}+kT_{D}\ln\left( \frac{E_{D}}{E_{A}}-1 \right)}.$$

For this general shape, we vary the TPC of the prey and predator relative each other (see methods), by shifting their TPCs in opposite directions along the temperature axis such that the $T^{*}$ values for each species are an equal distance from the mean environmental temperature. To allow for this shift, we fix $T_{D}$ and $E_{D}$ and manipulate $E_{A}$ separately for each species such that $T^{*}$ for each species takes the desired value. Specific parameter choices are detailed in Table S1.

***S1.2 Temperature Regimes***

We subject species to four different temperature regimes, which we select for easy comparison across regimes. Using the names defined in the main text, the temperature $T(t)$ in each regime is as follows, where $t_{max}$ is the end of the simulation horizon.

Variability only:

$$T\left( t \right)=\bar{T}+\left( T_{max}-T_{min} \right)\left( 0.5-2\left| t-floor\left( t \right)-0.5 \right| \right)$$

Warming only:

$$T\left( t \right)=T_{min}+\left( T_{max}-T_{min} \right)\frac{t}{t_{max}}$$

Warming and constant variability:

$$T\left( t \right)=T_{min}+\left( T_{max}-T_{min} \right)\left( \frac{t}{t_{max}}+\left( 0.5-2\left| t-floor\left( t \right)-0.5 \right| \right) \right)$$

Warming and increasing variability:

$$T\left( t \right)=T_{min}+\left( T_{max}-T_{min} \right)\left( \frac{t}{t_{max}}+\left( 1+\frac{t}{t_{max}} \right)\left( 0.5-2\left| t-floor\left( t \right)-0.5 \right| \right) \right)$$

Variability with stochastic events:

$$T\left( t \right)=\bar{T}+\left( T_{max}-T_{min} \right)\left( 0.5-2\left| t-floor\left( t \right)-0.5 \right| \right)+\xi\left( t \right)$$

where $\xi$(t) is a temperature shift taking value 0.5 iff a heat shock is occurring during the year defined by the interval [floor(t), ceiling(t)], and 0 otherwise. In other words, temperature shifts upward 0.5 degrees K during years in which an “event” occurs. The event $\xi$(t) is assumed to follow a Markov process with transition probabilities defined as follows:

$$P\left( E\left( \tau\right)=0.5 | E\left( \tau-1 \right) \right)=\left\{ \begin{aligned} 0.5 if E\left( \tau-1 \right)=0.5 \\ 0.07 if E\left( \tau-1 \right)=0 \end{aligned}, \right.$$

where $\tau$ denotes an integral value of t. The shift and transition probabilities are chosen to correspond approximately to El Niño events in Chile (following Miller et al 2019 [3]).

Variability with stochastic Gaussian process noise:

$$T\left( t \right)=\bar{T}+\left( T_{max}-T_{min} \right)\left( 0.5-2\left| t-floor\left( t \right)-0.5 \right| \right)+ \xi(t)$$

where $\xi\left( t \right)$ is a Gaussian process with squared exponential covariance function, but with variance and length-scale that change through time to allow for temperature deviations to change in magnitude and autocorrelation in the future.

Figures plot results as a function of the amplitude of temperature variability or warming, defined by $T_{max}-T_{min}.$ We increase amplitude by raising $T_{max}$ and lowering $T_{min}$ by identical amounts; thus increases in amplitude amount to mean preserving spreads. Results use $t_{max}=2000$unless otherwise indicated.

***S1.3 Partial Derivatives***

As stated in the main text, in the absence of temperature effects, the Rosenzweig-Macarthur model has the following coexistence equilibrium:

$$x^{*}=\frac{m}{a(c-mh)},$$

$$y^{*}=\frac{r}{a}\left( 1-\frac{x^{*}}{K} \right)\left( ahx^{*}+1 \right)=\frac{r}{a}\left( 1-\frac{m}{aK(c-mh)} \right)\frac{c}{c-mh}.$$

which requires that $aK\left( c-mh \right)>m$. The long-run predator population is increasing in *r*, while its dependency on the attack rate *a* is:

$\frac{{\partial y}^{*}}{\partial a}=\frac{c}{c-mh}\left( \frac{2rm}{a^{3}K\left( c-mh \right)}-\frac{r}{a^{2}} \right)$.

Assuming $c>mh$, so that the predator population is viable, $\frac{{\partial y}^{*}}{\partial a}>0$ iff $a<\frac{2m}{K\left( c-mh \right)}$. Thus, marginal increases in attack rates lead to increases (decreases) in equilibrium predator population levels for sufficiently low (high) starting attack rates. Because temperature variability and the relationship between TPCs and mean temperature can affect average attack rates, these partial derivatives can inform how temperature variability is likely to influence long-run predator population levels.

When a predator and its prey have the same TPCs, meaning that prey growth rate and predator attack rate respond identically to changing temperatures, only the effects of temperature on the attack rate matter. As a result, the prey is favored. Specifically, as seen in the deterministic coexistence equilibria, long-run prey levels are independent of $r$, while intrinsic growth only affects the long-run predator population through the ratio $r/a$. If the proportional effects of temperature on *r* and *a* are identical, which occurs if the TPCs for the species coincide, then the effects of temperature on equilibrium population levels will depend entirely on how temperature affects the attack rate. Temperature variability will lead to a lower average attack rate (and hence prey consumption) than if temperature remained at the optimum temperature for the predator, thus leading to lower long-run population levels for the predator, and higher levels for the prey.

***S1.4 Extension to multiple prey species***
When multiple prey species with different temperature affinities are available to the predator, different growth rates of those prey intuitively may reduce the likelihood that the predator population will collapse. To investigate this possibility, we generalize the predator population dynamics to account for multiple prey using a multispecies Holling type II response:

$$f\left( x\left( t \right), y\left( t \right), T\left( t \right) \right) =\frac{\sum_{j} a_{j}\left( T\left( t \right) \right)x_{j}\left( t \right)}{1+ \sum_{j} a_{j}\left( T\left( t \right) \right)h_{j}x_{j}\left( t \right)}.$$

Here, $j$ indexes the prey species. To illustrate the effects of multiple prey in the simplest setting, we focus on two prey species and assume that both $a_{j}\left( T\left( t \right) \right)$ and $h_{j}$ are invariant across species. To facilitate comparison with the single prey case in the main text, we conduct a series of experiments in which we shift the first prey TPC and the predator TPC in opposite directions as before, but fix the second prey TPC at the environmental mean. This keeps the degrees of freedom of our experiment fixed but permits us to illustrate how additional prey may dampen the threat of collapse. Results are depicted in Figure S5 and discussed in the main text.

***S1.5 Stochastic temperature shocks***

For the sake of exposition and analysis, the main text focuses on deterministic temperature changes in the form of seasonal variation, warming, or both. Here we briefly examine whether the patterns we identify also hold under stochastic temperature variation. We examine two types of stochastic temperature changes, as detailed above in Section S1.2. First, we examine major “events” modeled after El Niño; when such an event occurs, the temperature shifts up during a given year. Second, we examine smoother stochastic variability in the form of a Gaussian process with squared exponential covariance function. In both cases, we add these stochastic temperature deviations on top of our baseline scenario of oscillating (seasonal) temperatures. Doing so allows us to focus solely on the two types of variability rather than the combination of variability and warming.

For modest amounts of both types of stochasticity we examine, the addition of random variation in temperature does not change our qualitative conclusions (Figure S6), and follows intuitive patterns. For the modest levels of stochasticity in Figure S6, stochastic temperature changes knock population trajectories out of asymptotic convergence to equilibrium levels, but do not qualitatively change the long-run behavior. Other forms of stochasticity can, of course, have more dramatic effects. For example, stronger autocorrelation in temperature deviations can qualitatively change the long-run population dynamics. In the limit, a stochastic event with a continuation probability of 1 would correspond to a permanent shift of temperature, which could drive the predator population to low levels or extinction (Figure S7). More dramatic and unexpected changes are likely possible in a model in which temperature could influence mortality rates (large scale die-offs) or more mechanistic processes, but those are outside the scope of the Rosenzweig-Macarthur model on which we build.

**SI Tables**

**Table S1: Summary of model parameterization**

|  | Parameter | | Value | Notes |
| --- | --- | --- | --- | --- |
| Thermal performance curves | k | 8.62E-5 | | Boltzmann’s constant |
|  | $T^{*}$ | Varies by simulation. | | See methods |
|  | $T_{ref}$ | $T^{*}$ | | Setting $T_{ref}=T^{*}$ means $p(T_{ref})$ is the maximal rate, simplifying interpretation |
|  | $T_{D}$ | $T^{*}+1$ | |  |
|  | $E_{D}$ | 20 | |  |
|  | $E_{A}$ | $\frac{E_{D}}{1+e^{\frac{kE_{D}T_{D}^{2}}{T^{*}-1}}}$ | | Value ensures the selected $T^{*}$ is where the TPC maximum occurs. |
|  | $r\left( T_{ref} \right)$ | 0.6 | |  |
|  | $a\left( T_{ref} \right)$ | 0.3 for most results; 0.5 in panel c of Figs 2, S3, S4, S5 | |  |
| Temperature regimes | $T_{max},T_{min}$ | Varies by simulation. | | See section S1.2 |
| Other model parameters | K | 20 | |  |
|  | h | 0.3 | |  |
|  | m | 0.2 | |  |
|  | $c$ | 0.3 for most results; 0.1 in panel b of Figs 2, S3, S4, S5 | |  |

**SI Figures**

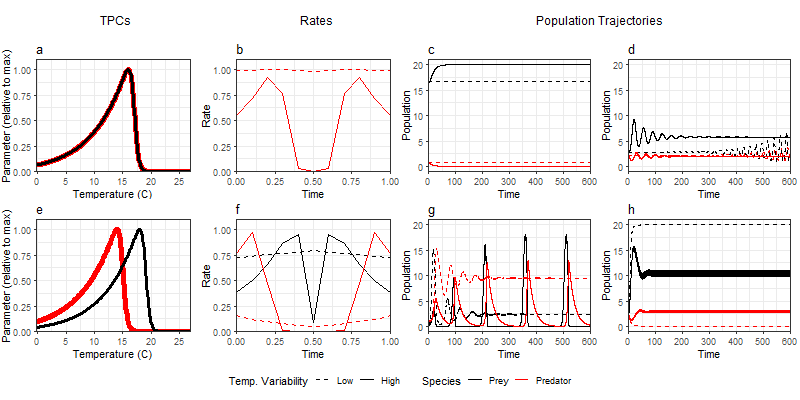
**Figure S1: Effects of temperature changes on population levels under different TPC configurations.**
Potential impacts of temperature variability on the predator (black) and prey (red) populations for low (dashed line) and high (solid line) temperature variability scenarios. Top row: thermal performance curves (TPCs) coincide (top row). Bottom row: Prey (predator) demographic rate is optimized at higher (lower) temperatures. First column: plot of TPCs relative to their maximum as a function of temperature. Second column: example one-period cycle of demographic rates that result from variable temperatures. Third and fourth columns: example population trajectories under different parameterizations.

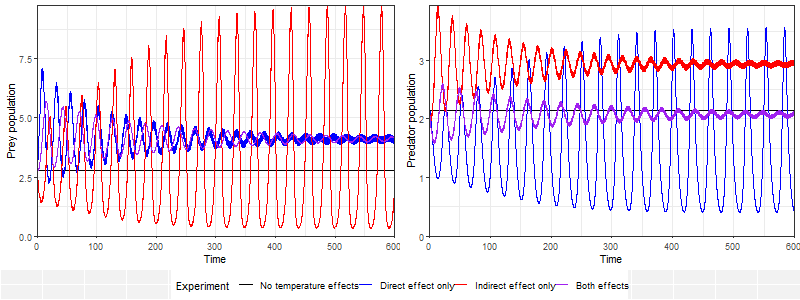
**Figure S2: Direct and indirect effects of temperature variability on population trajectories.**
Population trajectories for the prey (left) and predator (right) under different assumptions about which species is affected by temperature variability. Different lines examine cases when (1) neither species is affected by temperature (black), (2) the focal species is affected directly by temperature, but not the other (blue), (3) the other species is affected by temperature but not the focal species (red), and (4) both species are affected by temperature. Figures reflect TPCs that coincide for both species when a species is affected by temperature.

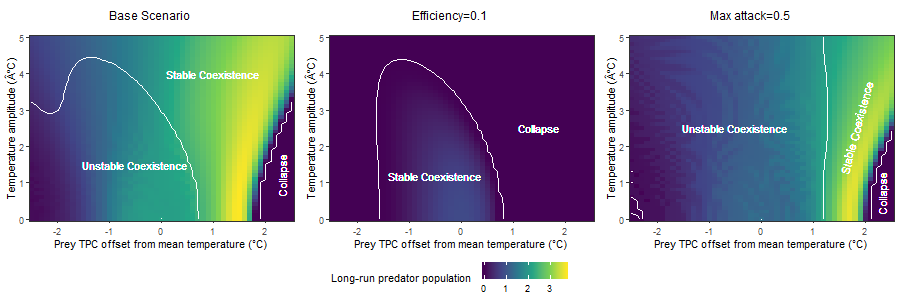

**Figure S3: Equilibrium types and levels using rates fixed at average levels.**
Long-run predator population levels (colors) and equilibrium type (labels, boundaries in white) as a function of prey TPC offset (x-axis; predator TPC offset is opposite that of prey) and amplitude of variability in temperature regime (y-axis) using *average* growth and attack rates ($\bar{r\left( T \right)} and \bar{y\left( T \right)}$) under the specified TPCs and temperature amplitude. Otherwise, population and temperature regime parameters parallel match those in Figure 2.

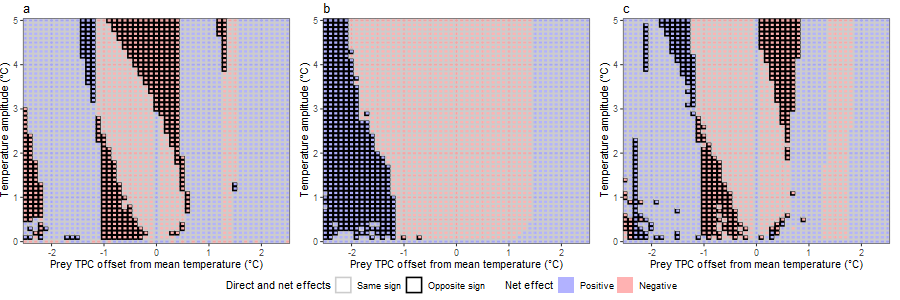

**Figure S4: Direct and net effects on predator population for other model parameterizations.**

Net effects (blue: positive; red: negative) of temperature variability on the long-run population levels for the predator, as a function of how far the prey TPC is offset relative to the mean temperature (x-axis) and the amplitude of temperature variability (y-axis). Outlines indicate whether the sign of those net effects matches (black outline) or does not match (light gray outline) the sign of the effect that would be predicted if only considering the effects of temperature variability on the predator, ignoring the temperature-dependence of its prey (‘direct effects’). Panels use different parameterizations matching those in Figure 2 panels a-c.

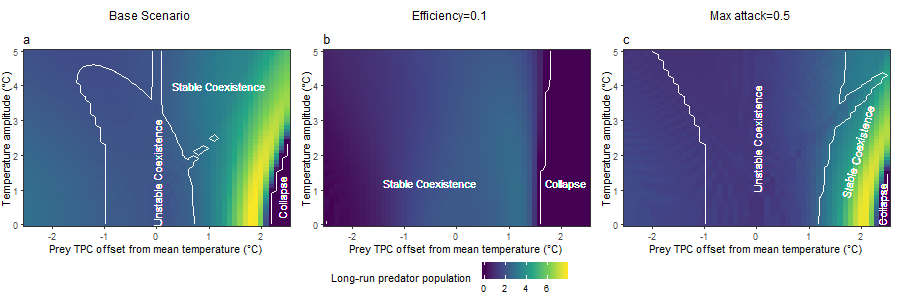

**Figure S5: Equilibrium types and levels with two available prey species.**

Effects of offset in the predator and prey TPCs (x-axis) and amplitude of temperature variability (y-axis) on predator population abundance (colors) and the type of equilibrium that arises: predator collapse or predator-prey coexistence, and whether the latter is stable or unstable (e.g., cycles or oscillatory behavior). Long-run predator population levels (colors) and equilibrium type (as labeled in the plots, with boundaries in white) as a function of first prey TPC offset (x-axis; predator TPC offset is opposite that of first prey) and amplitude of variability in temperature regime (y-axis). The second prey TPC is fixed with offset 0 (i.e., maximal growth occurs at the mean environmental temperature). Population and temperature regime parameters parallel match those in Figure 2; TPC parameters are listed in Table S1.

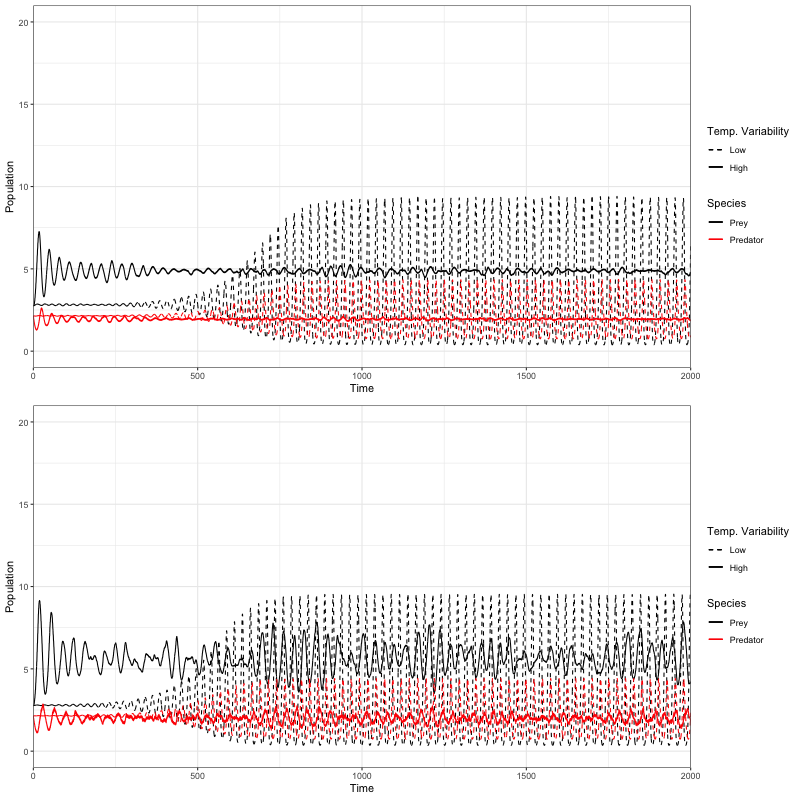

**Figure S6: Example population trajectories under stochastic temperature events.**
Potential impacts of temperature variability on the predator (black) and prey (red) populations for low (dashed line) and high (solid line) temperature variability scenarios. Predator and prey TPCs coincide with the mean environmental temperature, as in Fig S1 panel d. Top panel adds stochastic events following a Markov process each year; bottom panel adds Gaussian process noise.

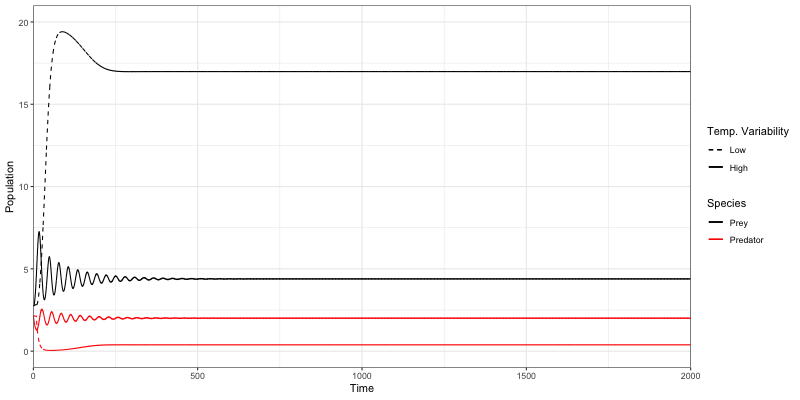

**Figure S7: Example population trajectories under highly autocorrelated stochastic temperature events.**
Potential impacts of temperature variability on the predator (black) and prey (red) populations for low (dashed line) and high (solid line) temperature variability scenarios. Predator and prey TPCs coincide with the mean environmental temperature, as in Fig S1 panel d. Stochastic events following a Markov process each year, but with a larger event effect (2 vs. 0.5 degrees in Figure S6) and with high probability of the event persisting once it occurs (1 vs. 0.5 in Figure S6).
